# Processing of Different Social Scales in the Human Brain

**DOI:** 10.1101/2022.08.01.502274

**Authors:** Moshe Roseman, Robin I. M. Dunbar, Shahar Arzy

## Abstract

For an individual to lead a healthy and fulfilling social life, it is essential to have relationships with multiple people who are at different levels of emotional closeness. Based on ethological, sociological and psychological evidence, social networks have been divided into five scales of emotional closeness, gradually increasing in size and decreasing in emotional proximity. Is this division also reflected in different brain processes? During functional MRI, participants compared their emotional closeness to different members of their social network. We examined the brain area that was differentially activated for levels of emotional closeness, and found that its vast majority (78%) showed preference for people who are closest to participants, including the temporoparietal junction, middle temporal gyrus, precuneus and dorsomedial prefrontal cortex. A different system, which includes the medial temporal lobe, retrosplenial cortex and ventromedial prefrontal cortex showed preference for all other social scales. Moreover, we found a significant correlation between brain responses to emotionally close people and smaller spaces (room, building) as well as between emotionally distant people and larger spaces (neighborhood, city). Finally, brain activity at the default mode network (DMN) was associated with social scale preference, such that its subnetwork DMN A, related to social processing, showed preference to closer social scales, while DMN C, related to spatiotemporal processing, showed preference to farther social scales. Our results show that the cognitive processing of a few intimately close people differs from the rest of the social network, emphasizing their crucial role in social life.

**Significance:** We divide the people in our lives according to levels of emotional closeness, called social scales, ranging from the few people who are the closest to us, in which we invest most of our social efforts and time (support clique), to the farthest level of ∼150 acquaintances. Here, we used neuroimaging to investigate the brain processing of different social scales. We found that the area of cortex dedicated to the support clique is much larger than that of all other scales and encompasses different brain regions. Interestingly, this division is similar to the one between processing of small and larger spaces, and processed by different subregions of the default mode network. Our study emphasizes the importance of close relationships in our social lives as found in the brain.

## Introduction

We encounter many people in our lives, but only a few of them will have a lasting impact on us. A large body of studies (e.g. [1]–[5]) suggests that, at any given time, each of us has only about 150 active connections with people, a figure which has become known as ‘Dunbar’s number’ [6]. Additionally, there are about 350 acquaintances which we know, but do not have meaningful relationships with. Together, these ∼500 people may be classified into five scales of personal closeness (social scales), successively identified as (1) support clique, (2) sympathy group, (3) affinity group, (4) active network, and (5) acquaintances. These scales run outward from the network owner, gradually increasing in size (with a constant ratio of ∼3) and decreasing in emotional proximity. This structure was identified in a variety of social contexts, including hunter– gatherer societies [3], [7], modern armies [8] and even mammalian social networks [9]. In addition, these layers have also been identified in a number of investigations of personal social networks using communication databases and social media (e.g. [5], [10]–[13]). Though the social scales have been investigated ethologically, sociologically and psychologically, not much is known about the brain processing underlying this classification. Do different cognitive mechanisms underlie the processing of different social scales?

Previous neuroimaging investigations of the effect of personal closeness on the processing of people compared people from one’s social network (i.e. all social scales together) [14]–[16] or a single social scale [17] to strangers. These studies identified differences in activity in specific brain regions, mainly the medial prefrontal and posterior cingulate cortices. More recent studies compared brain activity in response to people in one’s social network from different scales (rather than strangers) [18]–[21]. Specifically, Laurita, Hazan and Spreng [19] used a trait-judgment task to differentiate between romantic partners (scale 1) and other network members including parents (usually scale 1), close friends (scale 2) and acquaintances (scale 5). While processing of romantic partners involved a wide network of cortical regions, including the DMN components of the medial prefrontal cortex, the precuneus and the temporoparietal junction, all other individuals engaged only the parahippocampal cortex and the temporal pole. The authors suggested that the cognitive representations of close others are unique in content and use, chronically accessible and serve emotion-regulatory functions. In another study, differences between closer (scales 1-3) and less close (scales 4-5) people were based on activity in the medial prefrontal, inferior frontal and medial parietal cortices, and the temporoparietal junction [21]. Two other studies [22], [23] investigated people’s real world social networks (scales 1-5), and found that the temporoparietal junction, the posterior lateral temporal lobe and the retrosplenial cortex are involved in processing personal closeness. Yet, these studies did not systematically investigate reflections of the different social scales in brain activity, though such a reflection has been shown in between spatial and temporal scales [24], [25].

In an initial attempt to track the full sequence of social scales, Wlodarski & Dunbar [26] used functional magnetic resonance imaging (fMRI) to examine social networks of participants. Importantly, the focus of this study was not the differences between social scales but rather the differences between kin and non-kin. Nonetheless, in one analysis these authors compared brain activity during processing of different scales, although this analysis assumed a monotonic relationship between brain activity and social scale. As a result, this approach identified brain regions in which activity consistently increased (prefrontal, lingual and somatosensory cortices) or decreased (posterior cingulate and retrosplenial cortices) during the processing of more distant social scales. However, in other domains such as numerosity, space and time, neural activity exhibits a Gaussian-shaped rather than monotonic tuning curve [24], [25], [27], [28]. Furthermore, previous findings have implicated brain regions that displayed selectivity for one or more social scales, which do not follow this assumption of monotonicity [19]. Here, we tested the hypothesis that the five social scales are reflected in different patterns of brain activity.

## Results

### Bipartition of the scale-selective social cortex

Participants provided detailed descriptions of their own social networks, which enabled us to create individually tailored stimuli to engage each of their social scales separately. During fMRI scanning, participants were asked to compare their emotional distance to people across the five social scales (**Figure 1**). Only the parameter of social scale was manipulated, enabling us to look for differences in brain response for the different scales. To investigate social scale-selective activity, we identified voxels showing a significant difference in response to task performance at the different scales, then assigned the scale with maximal brain activity in each voxel. Strikingly, the vast majority of voxels showed a clear preference for the first social scale (Scale 1: 77.55%; Scale 2: 0.17%; Scale 3: 3.43%; Scale 4: 4.4%; Scale 5: 14.44%). In addition, a bipartition of the cortical scale-selective regions was apparent; while selectivity for scale 1 (support clique) was observed at the precuneus, temporoparietal junction, dorsomedial prefrontal cortex, middle temporal gyrus, supplementary motor area and precentral sulcus, separate regions in the ventromedial prefrontal, retrosplenial and parahippocampal cortices and in the hippocampus showed selectivity for scales 2-5 (sympathy group, affinity group, active network, acquaintances) (**Figure 2A**). Scale 2 was the least preferred social scale, minimally distributed within the medial prefrontal and retrosplenial cortices. Next, the averaged brain activity for each scale in each region was calculated and a hierarchical clustering analysis was performed (**Figure 2B**). The analysis validated the separation between scale 1 and scales 2-5 and the cortical bipartition between regions that showed preference for either group. Similar results were observed when choosing preferred scales by fitting a Gaussian function to the beta value graphs at each voxel and locating its peak (**Figure S3**).

**Figure 1.**
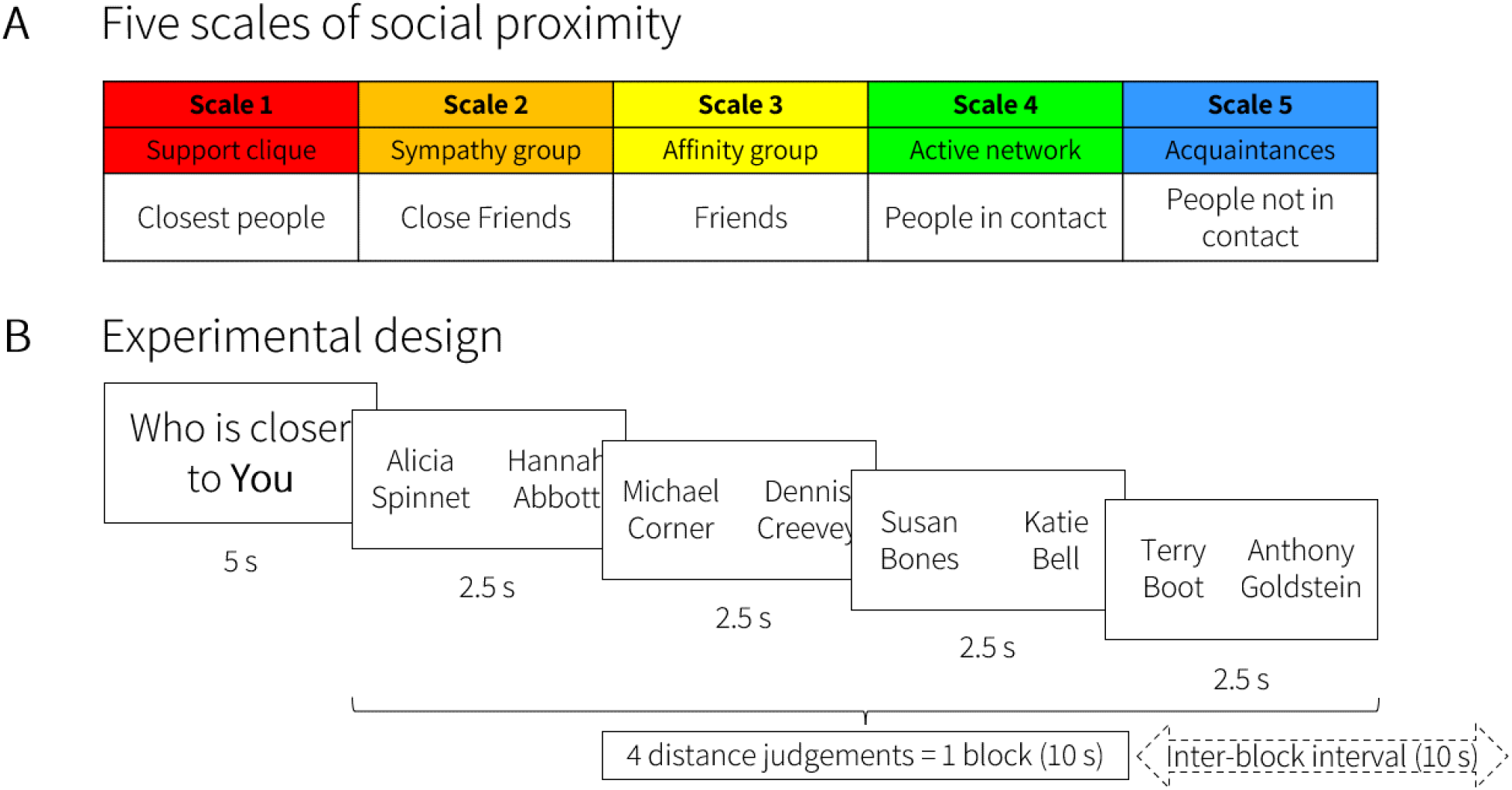
Experimental procedure. (**A**) Participants sorted members of their social network into five scales of social proximity, from closest friends to people with whom no contact is maintained. Participants were exposed only to the scale descriptions (bottom row) and not to the numbers or titles of the scales. (**B**) fMRI task. In each block, participants performed four social proximity comparisons, consisting of eight people from the same social scale.

**Figure 2.**
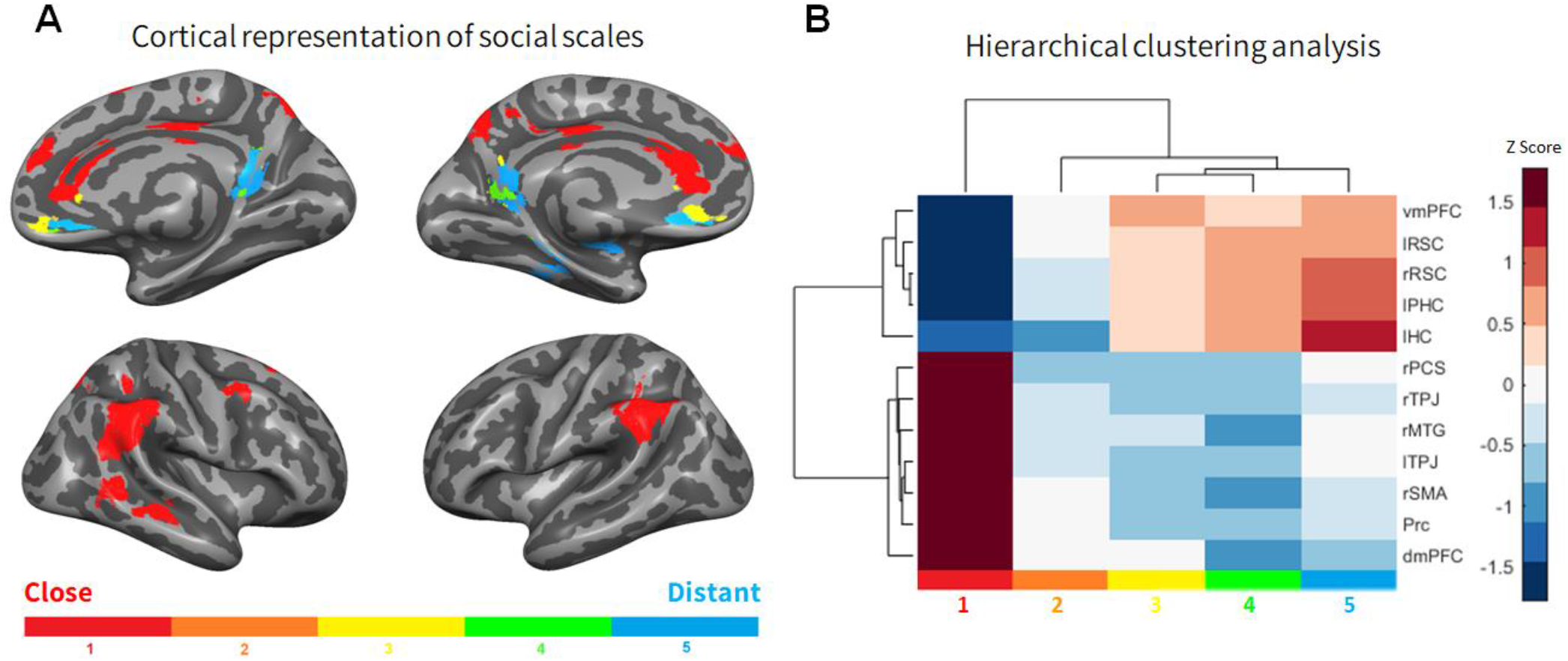
Social scale-selective activity shows a distinction between scale 1 (support clique) and more distant scales. (**A**) Scale-selective voxels identified by ANOVA across beta values (p<0.01, FDR-corrected for multiple comparisons, cluster volume threshold = 1000 mm^3^) and colored according to the scale of maximal activity across all participants. Note the dominance of scale 1 in the lateral cortical wall as well as in the dorsal aspect of the medial wall. (**B**) Hierarchical clustering analysis shows a bipartition between scale-selective brain regions that are tuned to scale 1 and scales 3-5, with scale 2 clustered closer to scales 3-5. (l, left; r, right; dmPFC, dorsomedial prefrontal cortex; HC, hippocampus; MTG, middle temporal gyrus; PCS, precentral sulcus; PHC, parahippocampal cortex; Prc, precuneus; RSC, retrosplenial cortex; SMA, supplementary motor area; TPJ, temporoparietal junction; vmPFC, ventromedial prefrontal cortex).

### Overlap between scales of different cognitive domains

Previous work showed that spatiotemporal and social functions rely on similar brain networks [18], [29]–[34] and that spatiotemporal processing is scale selective [24], [25]. To better understand the relations between the social and spatiotemporal functional networks and scales of proximity, we performed a voxel-wise comparison between brain representation of social scales as identified here and brain representation of spatial scales (room, building, neighborhood, city, country, continent), identified previously in a different cohort [24]. Small spatial scales (room, building) showed the largest overlap with the closest social scale (scale 1, or the support clique); medium spatial scales (neighborhood, city) showed the largest overlap with the most distant social scales (scales 4-5); large spatial scales did not show notable overlap with any of the social scales (**Figure 3**).

**Figure 3.**
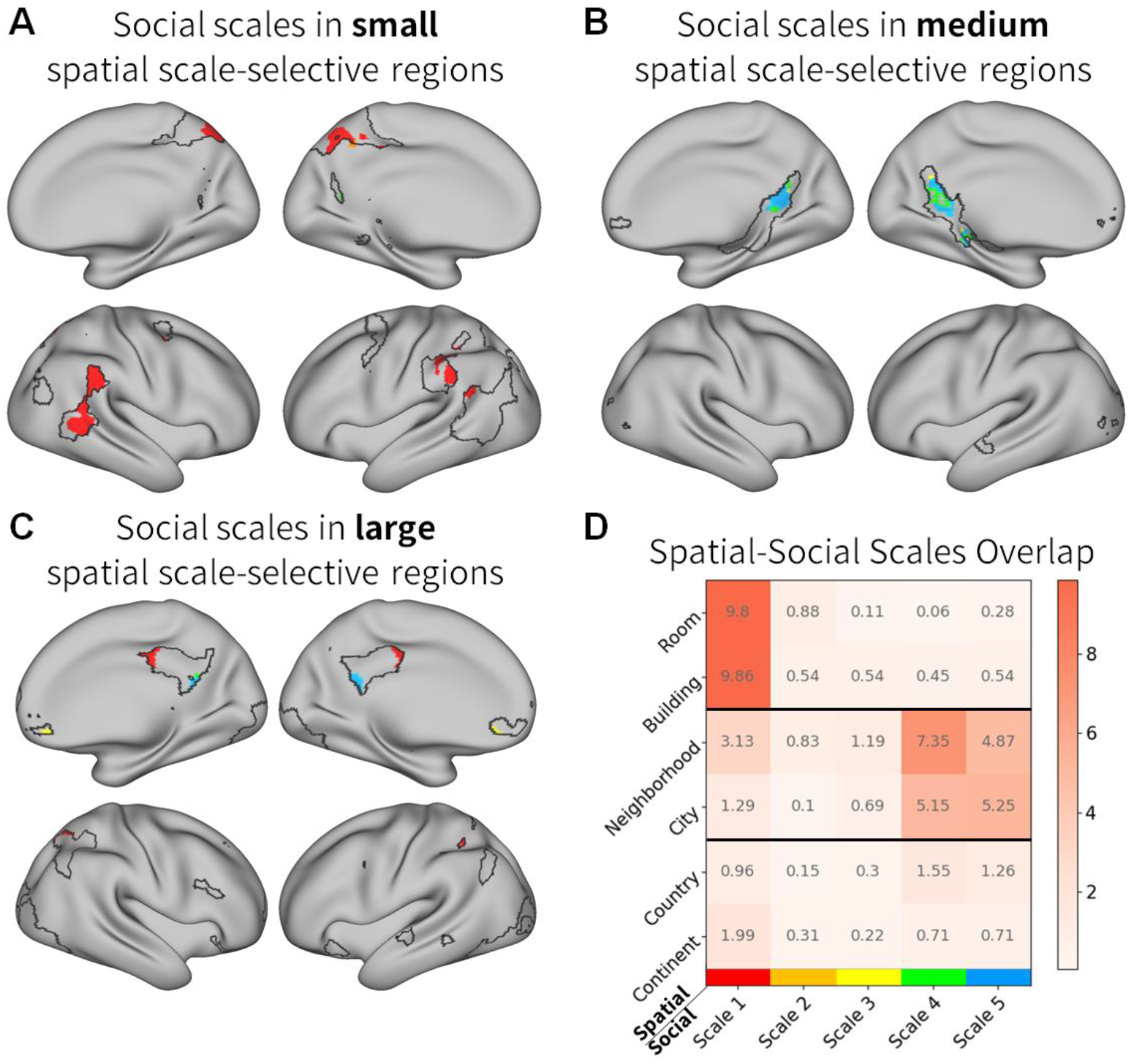
Spatial-Social scales overlap. (**A-C**) Social scale-selective activity in regions that also showed preference to small (room, building), medium (neighborhood, city) or large (country, continent) spatial scales. Spatial scale-selective ROIs (taken from [24]) are outlined in black. In these ROIs, voxels that showed significant selectivity to social scale (see Methods) are colored according to the scale with maximal activity. (**D**) Quantification of overlap between spatial and social scales. Values represent percentages from spatial scale-selective regions that showed selectivity to each social scale. While social scale 1 mainly overlapped with small spatial scales, social scales 4-5 mainly overlapped with medium spatial scales.

The relations between social and spatial scales are also reflected in their overlap with core regions of spatial and social cognition. Interestingly, when comparing social scale-selective regions to two partially overlapping domains of the brain’s orientation system, representing core spatial and social regions, closer social scales overlapped with core social regions, while more distant social scales overlapped with core spatial regions (**Figure S4**).

### Subnetworks of the default mode network exhibit different patterns of social scale selectivity

Previous work has decomposed the default mode network (DMN), a cortical network related to internal mentation and self-referenced activities, into subcomponents (DMN A, B and C), and associated DMN A and DMN C with social and spatiotemporal functions, respectively ([35], [36], see also [37]). To understand the relations between social scales and these cortical networks, we compared the anatomical distribution of the social scales to a parcellation of the human brain into seven cortical resting-state fMRI networks, as identified in data from 1000 participants [35]. The DMN showed the highest social-scale selectivity (**Figure 4A**). Examination of the relations between the social scales and the DMN subnetworks found that DMN A and DMN B showed significantly higher preference for scale 1 (support clique), while DMN C showed preference to the more distant scales 4-5 (**Figure 4B,C**).

**Figure 4.**
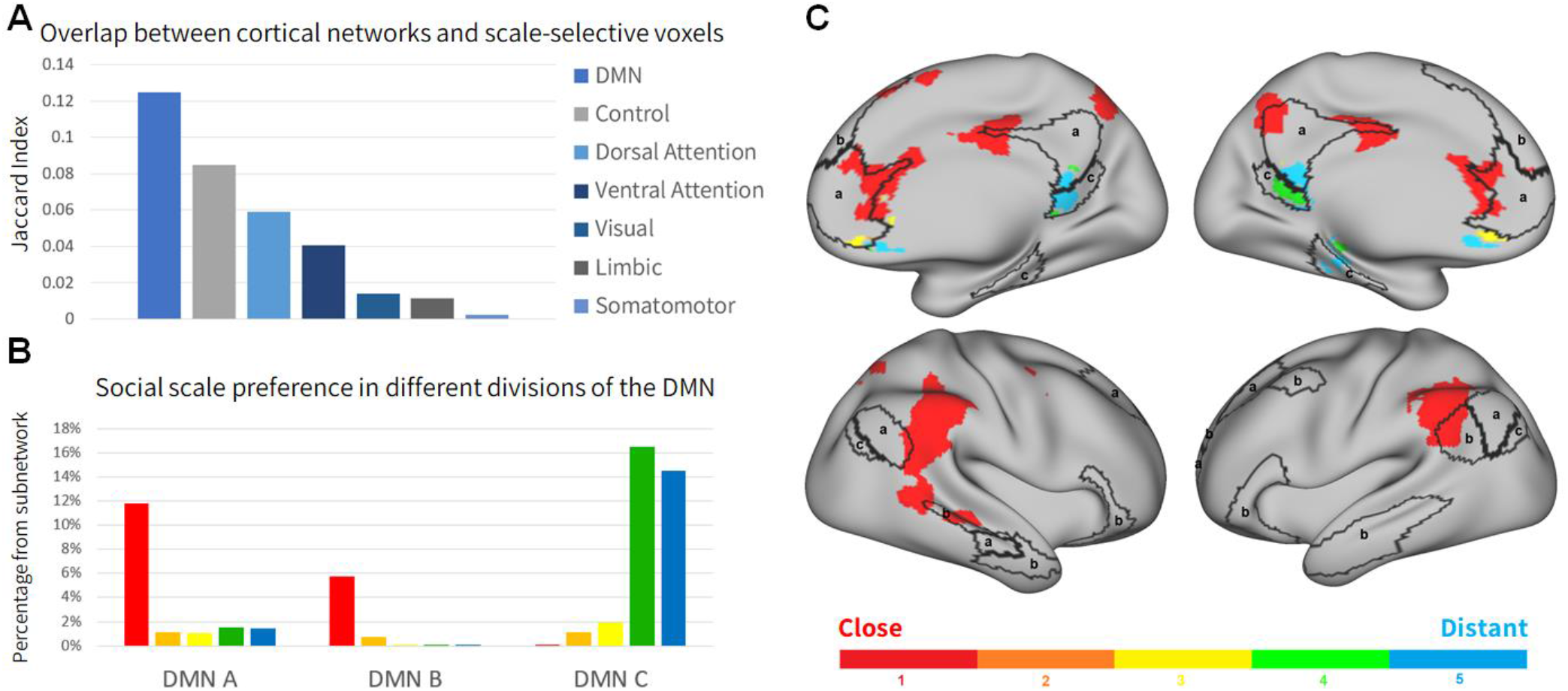
Social-scale selectivity of different cortical networks. (**A**) Overlap between 7 cortical networks [35] and scale-selective voxels. The DMN exhibits the largest overlap to scale-selective voxels. (**B**) Distribution of social scales within subdivision of the DMN (based on a parcellation into 17 cortical networks [35]). Percentages represent the proportion of voxels in each subnetwork that showed preference to that scale. (**C**) Cortical distribution of social scales with respect to subdivision of the DMN. Scale-selective voxels are identified and colored as in **Figure 2**. DMN regions are outlined in black and marked according to subdivision. While scale 1 is the dominant scale in DMN A and B, scales 4-5 are dominant in DMN C.

## Discussion

Examining how humans process members of their social network in different scales of personal closeness during fMRI scanning revealed several substantial novel findings. Firstly, a large cortical area differentially responded to the scales of one’s social network. Secondly, the vast majority of this area (78%) was selectively active for people in the closest scale; this area was anatomically segregated from regions that were selective for all other social scales. Thirdly, out of all the cortical networks, the DMN exhibited the greatest social scale preference; specifically, DMN A and B showed preference for the support clique while DMN C showed preference for distant social scales. Finally, social and spatial scales in the same proximity activated similar brain regions. This distinction between a small inner core and a larger outer core is also characteristic of the most social primates (cercopithecine monkeys and apes) [38]–[40], and seems to involve essentially the same brain regions as in humans [41], [42]. The fact that the inner and outer layers in social networks appear to be processed in different ways in separate parts of the brain offers a possible explanation for how large social groups are built up during the course of primate evolution. The present results suggest that this has been done by adding network layers using different cognitive mechanisms for successive layers (see [39]), thereby creating the fractal pattern characteristic of these species social groups [43].

The vast cortical surface dedicated to the closest social scale may reflect the significance of this scale to one’s social world. The closest social scale has been termed the support clique [1], since it encompasses the few closest people who are most likely to provide emotional and instrumental support in time of need [1], [3], [6], [44], [45]. The vast majority of one’s social cognitive efforts are dedicated to individuals in the support clique, which may account for the very small size of this scale [46]. At the same time, social reward provided by people in this scale significantly surpasses reward by people in all other scales and has been shown to be crucial to emotional well-being, stress reduction and healthy social interaction [2], [6], [47]. The disproportional cognitive effort and social reward associated with the support clique may account for its extensive representation in the brain despite its small size. Examination of the scale preference distribution along the cortex revealed a pronounced separation between regions that are tuned to the support clique and regions that are tuned to distant social scales. What may have led to such a marked partition? One possibility is that assessment of the personal distance between oneself and others recruits different cognitive mechanisms according to their social scale: in remote scales, psychological distance may play a significant role, whereas in the support clique additional aspects relating to the specifics of the relationship between the subject and these individuals may feature more prominently. Indeed, when asked whether their strategy differed between close and distant individuals, 48% of the participants mentioned that it did differ; according to their reports, these participants typically relied more on their “gut feeling” for closer people and more on temporal distance or episodic memory for distant people. Assuming that “gut feeling” judgments are more rapid than judgements based on mnemonic or spatiotemporal processes, these different cognitive mechanisms may also be reflected in the differences in response times, which were significantly shorter for the support clique with respect to all other scales (**Figure S1C**). Similar findings were reported by Wlodarski & Dunbar [26] who found that, when making decisions about social traits that applied to individual members of their personal social networks, reaction times for alters in the innermost layer (identical to our ‘close’ layer) were faster than to individuals who lay in more distant layers. Support for this possibility may come also from the implicated brain regions. The temporoparietal junction, dorsomedial prefrontal cortex, middle temporal gyrus and the precuneus, regions that showed preference to processing support clique individuals, are consistently implicated in processing the self and mental states of others [16], [19], [48]–[52]. In contrast, preference to more distant social scales was observed in the medial temporal lobe and retrosplenial cortex, regions that are associated with assessment of distance from allocentric and egocentric reference points, respectively [29], [53], [54] as well as with memory processes [55]–[57].

Another explanation for the difference in cognitive processes between different social scales is suggested by the observed neuroanatomical similarity between social and spatial scales, specifically that between distant social scales (scale 4, scale 5) and medium spatial scales (neighborhood, city). In physical space, small scale environments are represented in a precise manner, heavily influenced by visual perception, while large scale environments tend to evoke a more flexible representation that relies on map-like constructs [29], [58], [59]. Embedding information in a cognitive map and employing it during decision making were shown to involve the hippocampus, retrosplenial and ventromedial prefrontal cortices [53], [60]–[64], as was processing of large spatial scales [24]. Similarly, social tasks that require the embedding of relational properties such as power, affiliation, popularity and competence activate the cognitive map system [22], [65]– [67]. Accordingly, this system was postulated to help assimilate social knowledge in order to flexibly make social comparisons [34]. Our results indeed locate the distant social scales in these exact regions, which implies that the processing of distant social scales relies on cognitive maps. It should be noted that no social scale correlated with the largest spatial scales (country, continent). It is possible that the associated brain regions will show preference to even more distant people, such as people that one remembers only vaguely, or alternatively people that one knows but not vice versa (e.g., celebrities). Further experiments are needed to test these speculations.

The parcellation of the cortex to networks of functional connectivity may provide another explanation for the observed bipartition. Social distance judgment significantly involved all main hubs of the DMN, a cortical network known for its activity during self-referential and internal mentation [37], [68], [69]. While the distribution of support clique preference mainly follows DMN A, the distribution of distant social scales mainly activated DMN C. DMN A and DMN C were identified in a large group-level analysis and correlated with the “core” and “medial temporal” subnetworks, respectively, as defined in another parcellation of the DMN [35], [36]. Though the functional difference between these subnetworks is not fully understood, tasks involving theory of mind consistently involve the core subnetwork, while the medial temporal subnetwork mainly supports mnemonic and spatiotemporal processes [36]. Likewise, when DMN parcellation into two subnetworks is defined in individual subjects, one subnetwork is consistently implicated in theory of mind while the other one is implicated in episodic memory [70], [71]. Our results corroborate and extend this by demonstrating a differential recruitment of two DMN subnetworks according to social scale. The overlap between the support clique regions and the DMN subnetwork implicated in theory of mind may be reflected in the ease with which we assess the mental states of people who are close to us [72], [73]. Concurrently, the association of distant social scales with the medial temporal subnetwork may reflect the mnemonic contributions to distance assessment.

Finally, the cortical distribution of social scales in relation to the general activity during social distance processing (**Figure S2**) may also suggest an organizing principle for the cognitive mechanisms behind social distance judgments according to scale. The ventral portion of the precuneus was significantly engaged during social distance assessment regardless of scale. While the adjacent dorsal precuneus showed selectivity to the closest scale, adjacent ventral regions including the retrosplenial cortex showed selectivity to farther scales. Interestingly, a similar division of the medial parietal lobe into dorsal and ventral parts, respectively processing small and large scales, may also be found in the spatial and temporal domain [24], [25]. Our results demonstrate a similar pattern in the medial prefrontal cortex, though the task vs. control contrast mainly overlapped its ventral portion. This pattern implies a dorsal-ventral gradient along the medial wall, upon which dorsal and ventral regions are recruited as task demands require processing of close and distant scales, respectively. A similar gradient was discovered in an analysis of global connectivity based on a large dataset of brain activity at rest [74], [75]. While the first gradient (i.e., the gradient that explained the most variance) situated the regions implicated by our task vs. control contrast in one end of the spectrum and surrounding dorsal and ventral regions in the other end, the third gradient was spread along the dorsal-ventral axis of the medial cortical surface. This pattern is also associated with gradual changes in hippocampal connectivity, with greater concentration of fornix fibers in more ventral regions (consistent with the implication of the hippocampus in processing of distant scales), and in the connectivity patterns of large-scale networks [76].

Inevitably, our study has limitations. Since people are different in their social attributes, the definitions of each social scale may differ between participants. For example, one participant may consider the frequency of contact to be a major factor in identifying close friends, while for another, this factor may hold little significance. However, this general measure of psychological distance has proven itself reliable in many studies [6], [21], [32], [72]. To obtain valid representations of participants’ social networks, we did not provide exact definitions for each social scale but instead allowed them to choose the definitions that best fit their network. Inconsistencies in the social scale definitions between participants may thus introduce error variance. It is possible that some of the intertwining between the anatomical distribution of social scales are a result of this. However, the major separation between the support clique and the other scales is robust and in line with previous findings as described above. Other consequences of the flexibility in social network structure are the combination of kin and non-kin, and the potential difference between individual scales in terms of age, group affiliation or other factors. This may be important due to the potential difference in social information processing between friends and family [26], and should be investigated in future studies. Finally, the comparison between social and spatial scales was performed using two different cohorts. While these cohorts were similar in terms of age and occupation, identification of the social and spatial scales in the same cohort is required to validate our findings.

In conclusion, we have presented behavioral and neuroanatomical evidence to the separate cognitive mechanisms underlying processing of very close individuals and other members of our social network. Our results emphasize the unique role of close relationships in our social life and suggest that thought processes regarding these relationships may rely more on self-related processes that may be translated to “intuition” or “gut feeling”, while farther relationships are based more on cognitive mapping of world information. This study reflects the range of processes that make up social cognition, each contributing a distinctive element to the rich, multifaceted social world in which we live.

## Methods

### Participants

Twenty-six healthy participants (10 women, mean age = 26.1 ± 4.2 years) participated in the study. All participants provided written informed consent, and the study was approved by the ethical committee of the Hadassah Hebrew University Medical Center in conformity with the Declaration of Helsinki (2013).

### Task and procedure

A week before the experiment, participants were asked to disclose names of people with whom they have an active relationship, and to sort them into four categories representing the first four social scales. Participants were encouraged to use their phone contact list or social media to avoid forgetting anyone who may fall within these scales. Participants who failed to provide at least four names in each category were excluded from the experiment. For the fifth social scale, participants were asked to disclose the names of 30 acquaintances with whom they do not have an active relationship (**Figure 1**).

During the experiment, participants were presented with two names from their social network, both belonging to the same social scale, and asked to choose the name of the individual who was closer to them (**Figure 1**). Participants were instructed to indicate their response by pressing either the left or the right button. Stimuli were presented in a randomized block design. Before each block, the question “Who is closer to you?” appeared on the screen for 5 s. Next, four consecutive stimuli pairs were presented, each for 2.5 s. Importantly, all names in a single block belonged to the same social scale. Each block was followed by a fixation cross, resulting in an inter-block interval of 10 s. Participants were instructed to respond accurately but as fast as possible. The experiment consisted of five experimental runs for each participant, each run containing 20 blocks in a randomized order (four blocks for each of the five social scales). In total, participants performed 400 comparisons during the experiment. Participants’ responsiveness was generally good; all participants except one responded to >95% of the stimuli. One participant did not meet this criterion and was excluded from subsequent analyses. In addition, participants completed a lexical control task in a separate run, in which they viewed a target name followed by four name pairs taken from their experimental stimuli, and indicated which of the two names is closer to the target name in alphabetical lexical order. Stimuli were presented using the Presentation software (Version 18.3, Neurobehavioral Systems, Inc, Berkeley, CA, www.neurobs.com, RRID: SCR_002521). A training task using similar stimuli with names of fictional characters was delivered before the experiment. After completing the training, all participants reported that they understood the task. After the experiment, participants rated the level of difficulty for proximity judgements in each social scale. In addition, they were asked to describe the strategy used for determining responses for different social scales.

### MRI Acquisition

Participants were scanned in a 3-T Siemens Skyra MRI at the Edmond and Lily Safra Center neuroimaging unit. BOLD contrast was obtained with an EPI sequence (repetition time [TR] = 2500 msec; echo time [TE] = 30 msec; flip angle = 72°; field of view = 192 mm; matrix size = 96 × 96; functional voxel size = 2 × 2 × 2 mm; 60 slices, multi-band acceleration factor = 2, interleaved acquisition order; 164 TRs per run). In addition, T1-weighted high-resolution (1 × 1 × 1 mm, 160 slices) anatomical images were acquired for each participant using the magnetization prepared rapid gradient echo protocol (TR = 2300 msec, TE = 2.98 msec, flip angle = 9°, field of view = 256 mm).

### MRI Preprocessing

fMRI data were processed and analyzed using the BrainVoyager 20.6 software package (R. Goebel, Brain Innovation, Maastricht, The Netherlands, RRID:SCR_013057), Neuroelf v1.1 (www.neuroelf.net, RRID:SCR_014147), and in-house Matlab (Mathworks, version 2020a, RRID:SCR_001622) scripts. Pre-processing of functional scans included slice timing correction (cubic spline interpolation), 3D motion correction by realignment to the first run image (trilinear detection and sinc interpolation), high-pass filtering (up to two cycles), smoothing (full width at half maximum (FWHM) = 4 mm), exclusion of voxels below intensity values of 100, and co-registration to the anatomical T1 images. Anatomical brain images were corrected for signal inhomogeneity and skull-stripped. All images were subsequently normalized to Montreal Neurological Institute (MNI) space (3 × 3 × 3 mm functional resolution, trilinear interpolation).

### fMRI Analysis

#### Estimation of cortical responses to each social scale

A general linear model (GLM) analysis [77] was applied at each voxel, where predictors corresponded to the five social scales. Each modeled predictor included all experimental blocks at one social scale, where each block was modeled as a boxcar function encompassing the four distance comparisons. Predictors were convolved with a canonical hemodynamic response function, and the model was fitted to the BOLD time-course at each voxel. Motion parameters were added to the GLM to eliminate motion-related noise. In addition, white matter and CSF masks were manually extracted in BrainVoyager for each participant (intensity >150 for the white-matter mask and intensity <10 with a bounding box around the lateral ventricles for CSF), and the average signals from these masks were added to the GLM to eliminate potential noise sources. Data were corrected for serial correlations using the AR(2) model and transformed to units of percent signal change. Subsequently, a random-effects analysis was performed across all participants to obtain group-level beta values for each predictor.

#### Identification of clusters with social scale-sensitive activity

To identify clusters with differences in brain activity between social scales, single-factor repeated-measures ANOVA was applied in each voxel on the scale-specific predictors’ beta values, across all participants (FDR-corrected for multiple comparisons across voxels, p<0.01, cluster volume threshold = 1000 mm^3^). Beta values from identified voxels were used to determine selectivity to social scales in two methods: (1) beta values for each social scale were averaged across participants and the scale that demonstrated the maximal activity was selected; (2) a Gaussian function was fitted to the beta’s graph and the location of its peak was identified [24]. Gaussian fitting was performed for a vector of five beta values (for the five social scales) for each voxel of each participant, after its normalization by subtracting its minimum value. Fitting was performed using MATLAB, with bounds of 0 to 100 for amplitude, -100 to 100 for center, and 0 to 100 for width. In each voxel, peaks of Gaussian fits that passed a threshold of r^2^ > 0.8 were averaged across participants and rounded to yield the preferred social scale.

#### Assessment of social scale-selectivity distribution across large-scale resting-state networks

A previously published whole-brain parcellation into seven large-scale brain networks was used as a template for resting-state networks location [35]. The overlap between the distribution of scale-selective voxels and the seven cortical networks was calculated using their intersection over union (Jaccard index). To examine social-scale selectivity in subnetworks of the DMN, we used three resting-state networks that were identified as DMN components in a parcellation into 17 large-scale brain networks published along with the one mentioned above.

#### Comparison between social and spatial scale-selective regions

Overlap between social and spatial scale-selective regions was calculated using a previous fMRI study of spatial scales [24]. In that study, participants were shown a target location, then were asked to compare between two different locations and choose the one that is closer to the target. Importantly, locations occupied a physical space that represented one of six spatial scales (room, building, neighborhood, city, country, continent). Brain activity during processing of different spatial scales was compared in methods similar to those described here. Voxels that showed significant spatial scale preference were divided into three groups according to the preferred spatial scale (small - room or building; medium - neighborhood or city; large - country or continent). Overlap between spatial scale-selective and social scale-selective voxels was calculated for each spatial and social scale. Additionally, distribution of social scale-selective voxels was calculated in brain regions that were previously shown to be activated during a similar orientation task in the spatial and social domains, regardless of scale [32].

#### Comparison of activity to the lexical control task

Regressors for the lexical control were added to the scale predictors in the GLM analysis, and a new design matrix was computed for each participant. A group analysis (corrected for serial correlations, AR(2)) was performed in each voxel, and activity during the social proximity judgment task in all five social scales was contrasted with the activity during the lexical control task.

## Acknowledgments

MR is supported by the VATAT scholarship for Data Science doctoral students.

## Supplementary Results

### Behavioral Results

Analysis of participants’ ratings of task difficulty for each scale indicated no significant differences between scales (all ps >.05, one-way ANOVA, Tukey–Kramer post hoc test) (**Figure S1A**). When asked to describe their judgment strategy, nine participants (36%) reported using the same strategy for all social scales. Six participants (24%) reported relying more strongly on meeting frequency in distant social scales, and another six reported (24%) involving episodic memory for distant more than close social scales. Four participants (16%) did not provide an answer.

For each participant, the average response time in each scale was calculated (**Figure S1B**). Repeated-measures ANOVA was conducted on the averages using a linear mixed model with scale as a fixed factor and participant as a random factor. This analysis demonstrated a significant effect for scale (F(4, 80) = 25.55, MSE = 4807.54, p < 0.001). A Tukey–Kramer post hoc test showed a significant difference between Scale 1 and other scales (p = 0.0003), while no other consecutive scale pairs differed significantly.

### Brain activity during social proximity processing vs. a lexical control task

Comparison between brain activity during social proximity processing and a lexical control task implicated the precuneus, ventromedial prefrontal cortex, bilateral temporoparietal junction and the left superior temporal sulcus (**Figure S2**).

**Figure S1.**
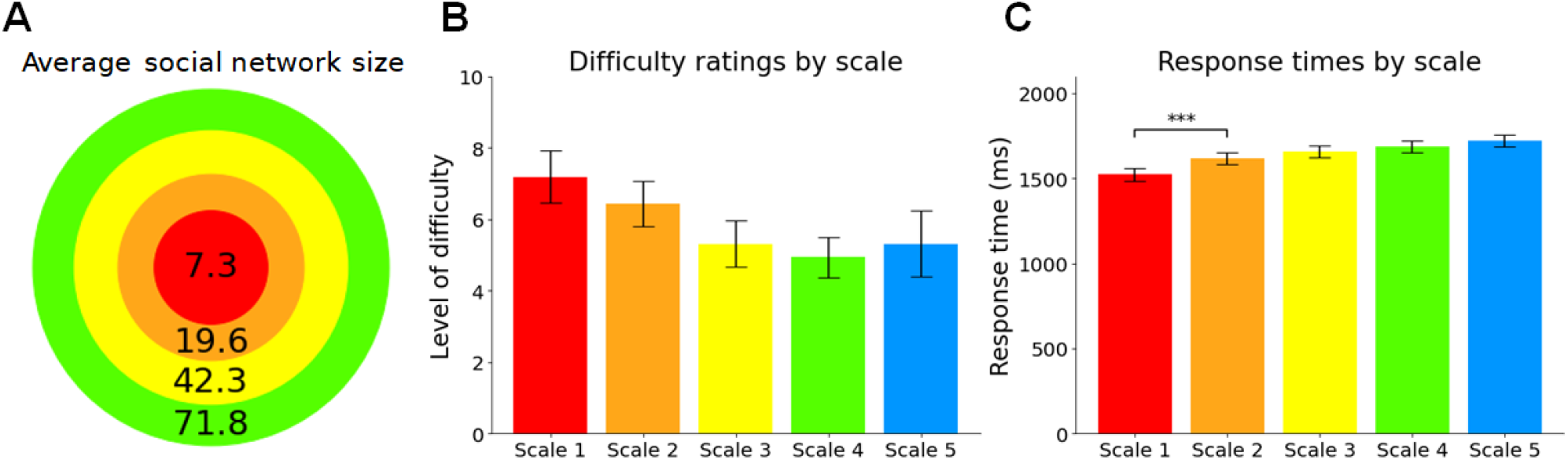
Behavioral results.(**A**) Reconstruction of Dunbar’s layer model [6] according to the average number of people in the social scales of the participants. Layers are hierarchically inclusive. Scale 5 (acquaintances) is omitted from the model as its size was fixed and dictated to the participants. (**B**) Level of difficulty as rated by the participants in a post-scan questionnaire. Error bars represent standard errors of the mean across participants. Differences between scales were not significant. (**C**) Response times (milliseconds). Error bars represent standard errors of the mean across participants. Significant differences were found between scale 1 (support clique) and more distant scales in repeated-measures ANOVA and Tukey–Kramer post hoc tests (*** p < 0.001).

**Figure S2.**
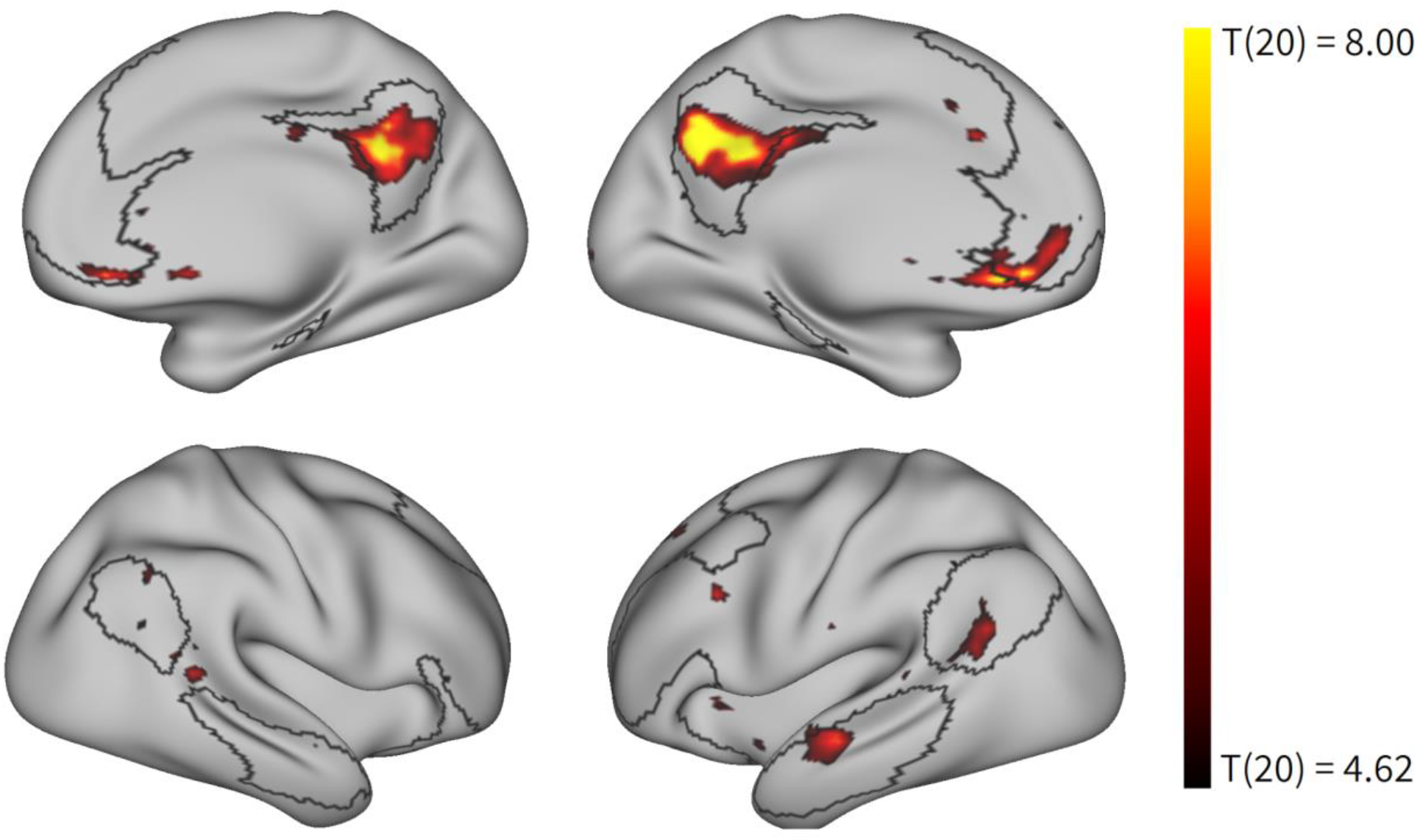
Task vs. Control. Colored voxels represent the group-level average of brain activity during the social task versus a lexical control task (p<0.01, FDR-corrected for multiple comparisons). This comparison implicated the precuneus, ventromedial prefrontal cortex, temporoparietal junction and the left superior temporal sulcus. DMN regions (as defined in [35]) are outlined in black.

**Figure S3.**
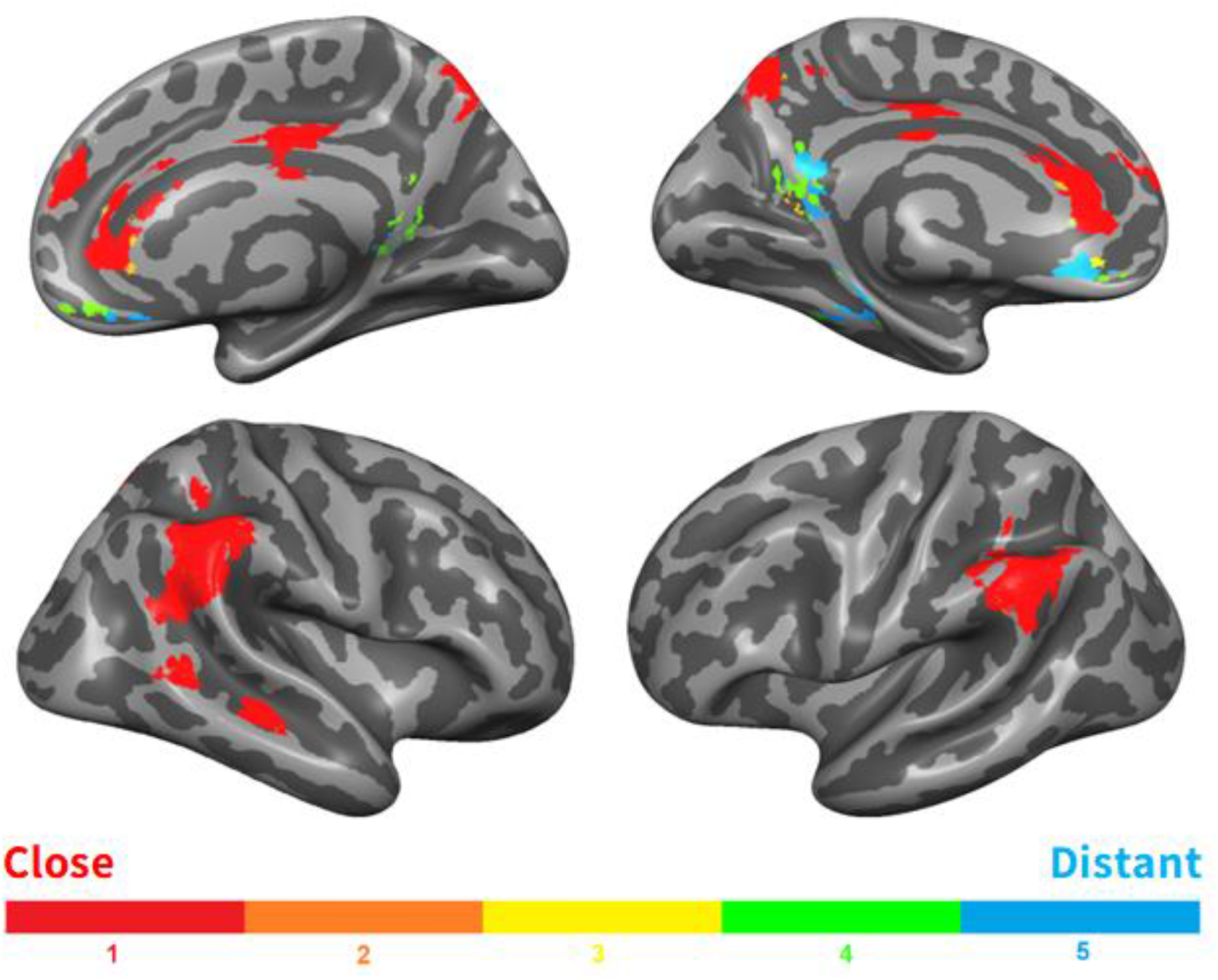
Scale-selective voxels were identified by ANOVA across beta values (p<0.01, FDR-corrected for multiple comparisons, cluster volume threshold = 1000 mm^3^) and colored according to the position of the peak of the fitted Gaussian (minimum r^2^ of fit = 0.8).

**Figure S4.**
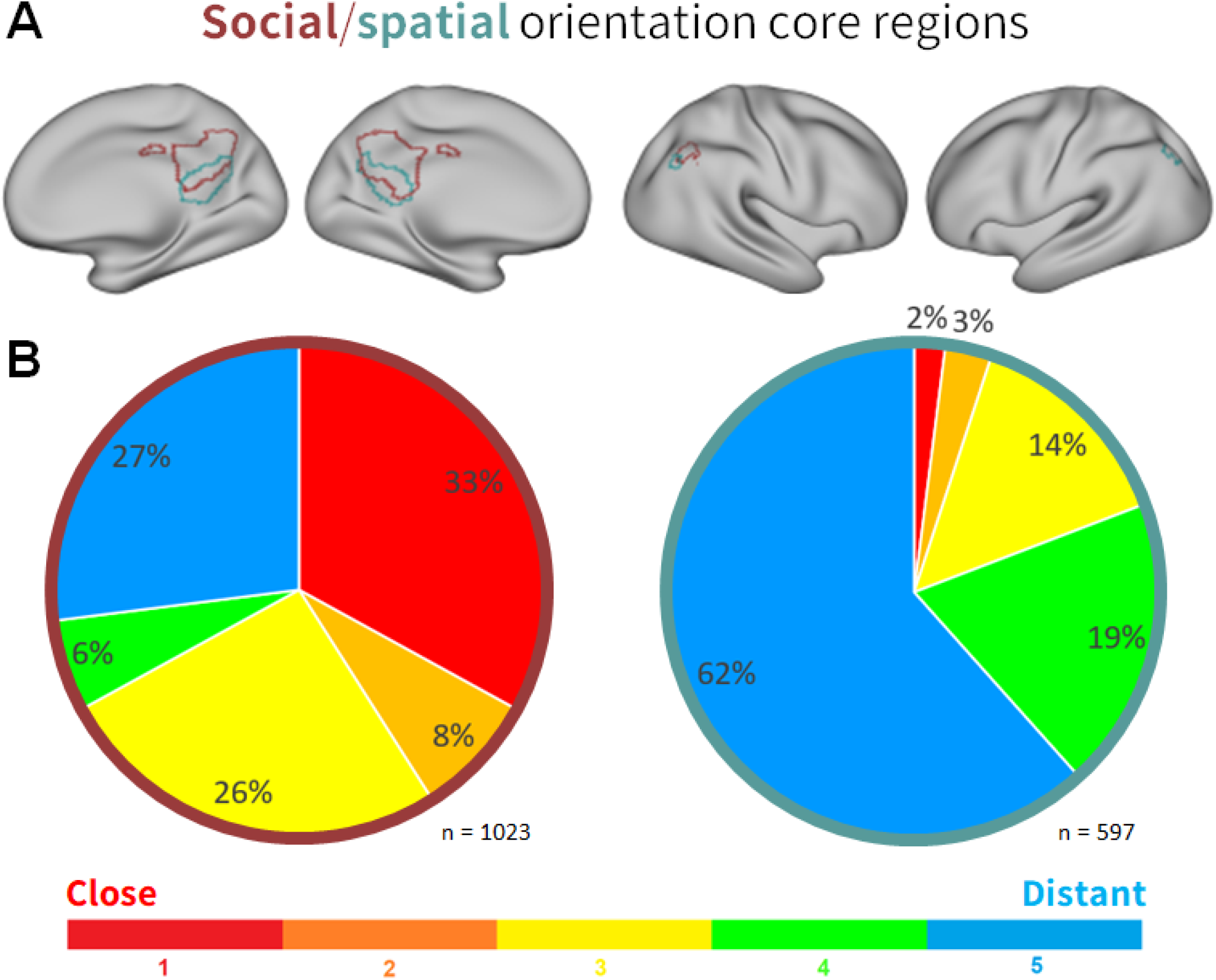
Social scale selectivity in orientation core regions. (**A**) Core regions for spatial (teal) and social (maroon) orientation were defined according to a previous experiment with similar methods that examined orientation in several cognitive domains regardless of scale [32]. (**B**) Distribution of social scales within social (left) and spatial (right) orientation core regions. Scales were assigned to voxels according to the maximal participant-averaged beta weights. Though there is overlap between the two domains, social orientation core regions were more tuned to close social scales than spatial orientation core regions are.

